# Tactile and Visual Spatial Frequency Perception Follows Optimal Integration but Is Not Affected by Spatial Proximity

**DOI:** 10.1101/2025.05.27.656312

**Authors:** Guandong Wang, David Alais

## Abstract

Spatial frequency is a fundamental feature in both visual and somatosensory perception, yet how these modalities integrate spatial frequency information remains unclear. This study investigates whether visuotactile spatial frequency perception follows the principles of Maximum Likelihood Estimation (MLE) and whether spatial proximity influences multisensory integration, using virtual reality (VR) and high-precision 3D-printed tactile stimuli. Experiment 1 found that the visuotactile integration of spatial frequency cues follows the MLE rule. However, Experiment 2 revealed that this integration is not affected by spatial proximity. These findings provide additional insights into the feature dependency of multisensory integration between vision and touch and highlight the potential for independent processing before integration, offering new perspectives on the mechanisms of spatial frequency processing in both visual and tactile modalities.

## Introduction

Spatial frequency, which describes how often surface structures change periodically in space, is a key aspect of human perception. In vision, spatial frequency perception enables the visual system to efficiently extract features from complex scenes by processing different spatial scales through dedicated frequency channels (Kauffmann et al., 2014; Lamb & Yund, 1996; Sachs et al., 1971). This function is supported by spatial frequency-selective neurons, which are found throughout the visual hierarchy, from ganglion cells (Kelly, 1975) to cortical neurons (De Valois et al., 1982; Foster et al., 1985). In the primary visual cortex (V1), the receptive fields of neurons can be modelled as Gabor filters, narrowly tuned to specific orientation and spatial frequency (Pollen & Ronner, 1983). These neurons respond optimally to stimuli matching their preferred orientation and spatial frequency. Through the combined population response of neurons with diverse tuning properties, V1 processes visual elements like contours and shapes across different frequency bandwidths (DeValois & DeValois, 1990).

In somatosensory perception, spatial frequency also plays a crucial role in sensory processing, as objects and surfaces encountered through touch exhibit both low spatial frequency features (such as shape) and high spatial frequency features (such as texture or roughness). Research on the receptive field properties of primary somatosensory neurons indicates that Gabor-like filtering of spatial variations in cutaneous afferent signals contributes to roughness perception (Hsiao et al., 1993). And a substantial amount of somatosensory neurons have been shown to be tuned to specific spatial frequencies (Bourgeon et al., 2016). These findings demonstrate that the neural basis and processing mechanisms of spatial frequency perception are highly similar between touch and vision.

In the natural environment, it is rare for our perceptual system to only depend on unisensory information. Instead, the brain typically receives multiple streams of information from different sensory modalities simultaneously. Therefore, it is crucial for brain to utilise the multisensory stream of information to enhance precision and resolve ambiguous perceptions. One prominent framework used to model multisensory integration is the Maximum Likelihood Estimation (MLE) model (Alais & Burr, 2019). The MLE model suggests that the brain combines sensory information in a statistically optimal manner, where the integrated multisensory percept is a weighted linear combination of cues from different modalities. The weight of each sensory input is proportional to its reliability, resulting in a more precise percept than individual unisensory estimates. This optimal cue combination has been validated across a wide range of behavioural studies, including visual-auditory (Alais & Burr, 2004) and visual-vestibular (Fetsch et al., 2010), etc. Specifically, MLE integration has been shown in various features between tactile and vision (Ernst & Banks, 2002; Helbig & Ernst, 2007b; Helbig et al., 2012). In addition, a series of studies highlight the close connection between vision and touch (Lunghi & Alais, 2013; Sathian et al., 1997; van der Groen et al., 2013; Wang & Alais, 2024). Hence, it is of interest to examine whether spatial frequency, as a fundamental aspect of both systems, integrates in an MLE fashion.

Despite the MLE model’s success in explaining many cases of multisensory integration, it sometimes fails to account for observed behavioural data. On the one hand, there were tasks that demonstrated a suboptimal integration between two modalities, such as audiovisual integration of temporal rate and spatial location cues (Arnold et al., 2019). On the other hand, there were also scenarios where a supralinear combination of multisensory information has been found, where the bimodal performance exceeded the statistical optimal linear combination of the two unimodal signals, like the integration of visual and tactile orientation information in rats (Nikbakht et al., 2018). There were also scenarios where the MLE prediction did not align well with the weightings of different modalities (Meijer et al., 2019). Moreover, in tactile-visual texture and roughness perception, where spatial frequency plays a key role, findings have been mixed: evidence has been found that supports both the two modalities being separate and independent (Guest & Spence, 2003; Roberts et al., 2024; Whitaker et al., 2008) or being an integrated system (Jones & O’Neil, 1985; Lederman et al., 1986) in processing the texture information. These findings together highlight the need for a deeper understanding of the rules governing multisensory integration. And specifically, in the context of spatial frequency perception.

Another key principle of multisensory integration is the spatial rule, which suggests that optimal integration occurs when cues from different modalities are spatially colocalised (Kadunce et al., 2001; Stein & Stanford, 2008). This principle stems from neurophysiological studies on the receptive field properties of single multisensory neurons (Kadunce et al., 2001; Stein et al., 1989; Stein & Stanford, 2008), and has been widely supported by evidence from a wide range of behavioural tasks (for reviews, see Spence (2013)). Specifically, it has been demonstrated in spatial distance judgement between vision and touch (Gepshtein et al., 2005). However, there is also substantial evidence from behavioural tasks that do not follow this pattern. Spence (2013) argues that the validity of the spatial rule is task-dependent; it tends to hold when spatial location information is relevant to the task, but often fails for tasks that do not require spatial information, such as identification and temporal judgment. The contradictory behavioural findings suggest that multisensory integration may not be confined to multisensory neurons that exhibit optimal responses to overlapping cues. Instead, there may be other concurrent pathways and alternative mechanisms that contribute to the complex interactions between multisensory cues.

As a localised feature, the judgment of spatial frequency is generally considered to be largely independent of spatial location in both vision and touch. However, in vision, the localised receptive fields of primary sensory neurons lead to spatial frequency processing being location-dependent (Williams et al., 1982). In touch, although spatial location is not explicitly required for the task, spatial attention has been shown to play a critical role in the processing of spatial frequency information (Sathian & Burton, 1991). Hence it remains an open question whether spatial frequency integration adheres to the spatial principle.

In this study, we aim to test whether MLE integration occurs between vision and touch in spatial frequency perception and to examine whether spatial frequency integration follows the spatial rule. By exploring the interplay between these modalities, we seek to better understand the conditions under which multisensory integration is successful.

## Methods

### Participants

A total of 51 participants were recruited for our study. Twenty-six participated in Experiment 1 and 25 participated in Experiment 2. All participants reported being right-hand dominant with no recent history of damage to the right index finger and no history of damage or diseases of the nervous system. This research was approved by the University of Sydney Human Research Ethics Committee (HREC 2021/048), and all methods were carried out in accordance with relevant guidelines and regulations.

Participants were first and second-year psychology students from the University of Sydney, recruited through the Sydney University Psychology SONApsych research participation system and were given course credit for their participation. Informed consent was obtained from all participants prior to the commencement of the experiment.

Six participants were excluded from Experiment 1 due to exceptionally low performance (<70% in at least one of the five conditions) to ensure sufficient data quality for reliable psychometric function fitting and parameter estimation Similar exclusion criteria were applied to Experiment 2: one participant was excluded for not completing the test due to personal reasons, and five were excluded due to exceptionally low performance.

## Apparatus and Stimulus

### Visual Stimulus

Visual stimuli used in the experiment were 3D models of circular disks with textured noise patterns, an example of the stimulus is shown in Figure 1a. The surface texture was created using a MATLAB toolbox (Wang et al., 2022), by passing white noise images of 500 × 500 pixels through 2D Gaussian band-pass filters. The surface textures of the disks were controlled by varying the peak frequencies of the band-pass filters, such that each stimulus disk had a different range of spatial frequency components. A 2D circular cosine ramp was then applied to the square noise pattern to create a smooth roll-off. The textured patterns were then converted to a 3D object using MATLAB and blender using the intensity of the grey-scale image as elevation.

**Figure 1.**
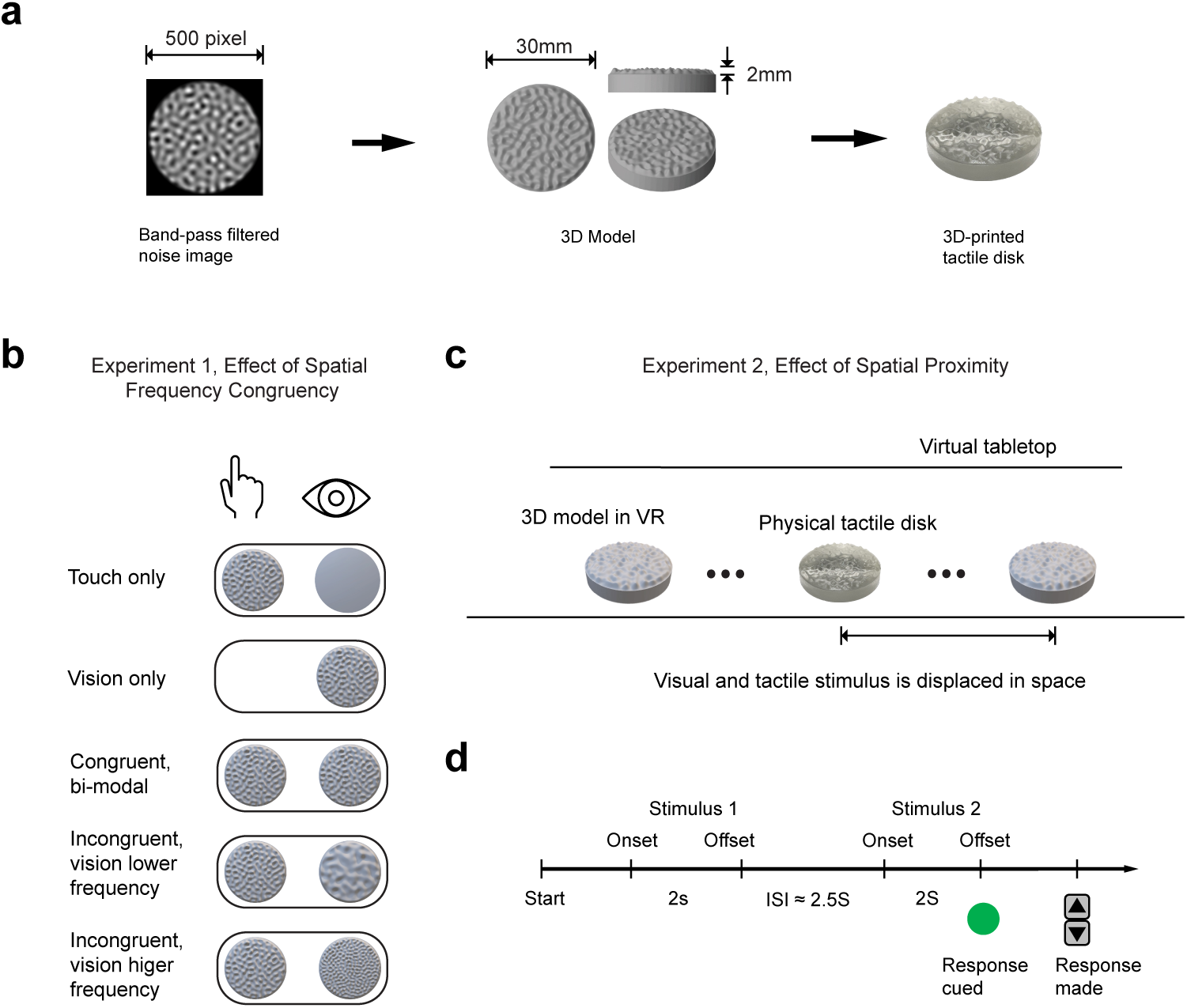
Experimental stimuli, conditions and procedures. *Note.* **a** From left to right: Band-pass filtered noise image, 3D model generated from the noise image used as the visual stimulus, and the corresponding 3D-printed tactile stimulus. **b.** Example of conditions for Experiment 1: Effect of Spatial Frequency Congruency. Five different conditions were tested in Experiment 1: two uni-modal conditions (touch-only and vision-only) and three bi-modal conditions (congruent bi-modal, incongruent with vision at a lower frequency, and incongruent with vision at a higher frequency). **c.** Demonstration of conditions in Experiment 2: Effect of Spatial Proximity. In Experiment 2, the visual stimulus was displaced horizontally from the physical stimulus to investigate the effect of spatial proximity on multisensory integration. The virtual device was replaced with a large (10 m wide) white virtual tabletop to minimize the potential for any spatial reference providing additional cues to the participants.

### VR Headset

The visual stimulus was presented to participants via a Vive Pro Eye VR headset, with 1440 × 1600 pixels per eye (2880 × 1600 pixels combined) resolution, 90 Hz refresh rate, and 110 degrees field of view. The headset was powered by the Dell XPS 8950 desktop equipped with an NVIDIA RTX 3070 graphic card.

### Tactile Stimulus

The tactile disks used in this experiment were fabricated using a Formlabs Form 3 industrial resin 3D printer. The author created the design for the 3D models and the disks were printed at the Sydney Manufacturing Hub. Formlabs Clear V4 photopolymer resin was used as the printing material, with both the layer height and lateral resolution set to 25 µm to ensure high surface texture quality.

### Tactile Device

The 3D-printed textured disks were presented to participants using an Arduino-controlled device. Five tactile disks were attached to a platform, which could be rotated by a stepper motor to position one disk underneath an aperture on top of the device. The tactile stimulus could then be delivered by raising and lowering the platform with a servo motor (Figure 2a). Additionally, the orientation of each disk could be adjusted by rotating another stepper motor. A Vive Tracker was fixed on top of the device, providing positional data that allowed the visual stimulus to be mapped onto the physical device in the VR environment.

**Figure 2.**
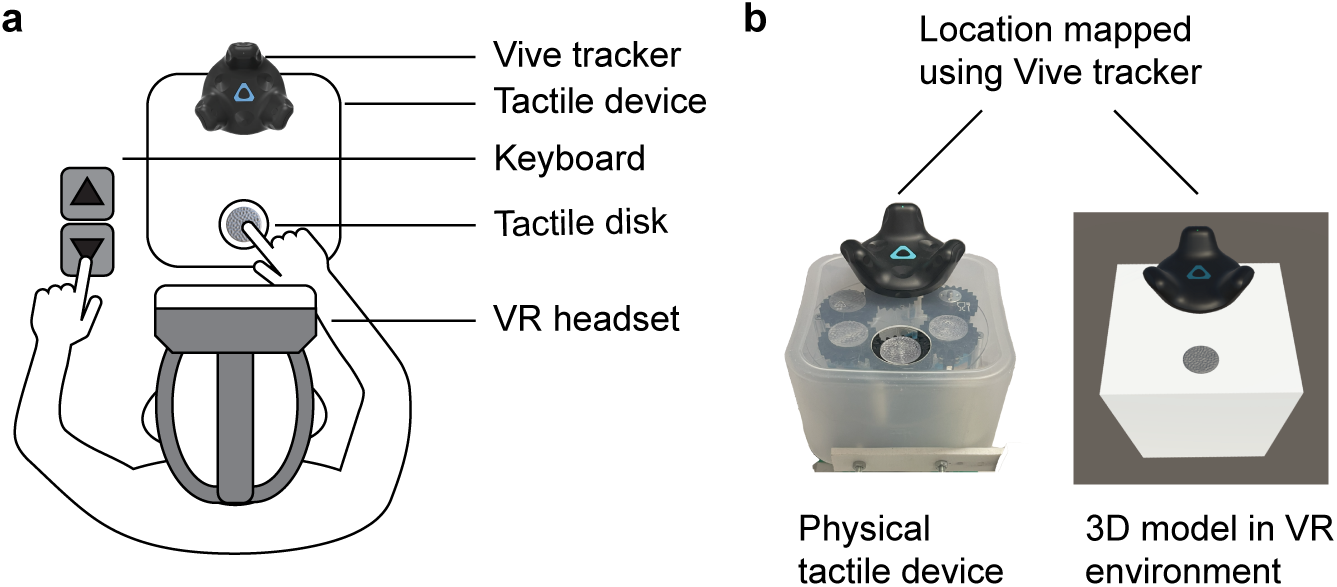
Experimental Setup and Apparatus. *Note.* **a.** Experimental Setup: Participants were seated in front of the tactile device while wearing a VR headset. Tactile stimuli were delivered using an Arduino-controlled device, while visual stimuli were presented through the VR headset. **b.** Left: Tactile device. Right: Device and stimulus presentation in VR. The physical device’s location was tracked using the Vive tracker, ensuring that the 3D model was rendered in the virtual environment at the same position. This alignment allowed the visual and tactile stimuli to be spatially co-localized.

### Experimental Environment and Visual-Tactile Spatial and Temporal Synchronisation

The experiment was scripted and designed using Unity 2021.3.16f1 and C#. 3D models of tactile disks were loaded into the virtual environment. The spatial colocalisation of the virtual (visual) and physical (tactile) stimuli was achieved using the Vive Tracker, which provided real-time coordinates of the physical device. The virtual device and stimuli were then rendered in real-time to match the physical location. The experimental process was controlled by C# scripts and Arduino code. Temporal synchronisation of the physical and virtual stimuli was achieved through serial communication between the computer and the Arduino device, where the visual stimulus was triggered to present once the sensor on the Arduino indicated that the tactile disk had reached the designated height (Figure 2b).

### Procedure

#### Experiment 1: Visuotactile Integration of Surface Spatial Frequency

Five different spatial frequency congruency conditions were tested in Experiment 1: two uni-modal conditions (vision-only and touch-only), and 3 bi-modal conditions (Congruent, Vision Lower Frequency and Vision Higher Frequency), as shown in Figure 1b.

The experiment followed a classic two-interval forced-choice (2IFC) paradigm. In the visual-only, tactile-only, and congruent bimodal conditions, the same set of five disks was used as the test stimulus in 2IFC trials, presented in vision, touch, or both, respectively. In the two incongruent bimodal conditions, the tactile stimulus was presented simultaneously with a visual stimulus of incongruent spatial frequency (one step lower or higher than the tactile stimulus; see Figure 1b). The reference stimulus in each 2IFC trial was a disk with a spatial frequency of 0.400 cycles/mm. The test and reference stimuli were presented sequentially in counterbalanced and randomised orders. Different conditions were also randomised and intermingled between trials. After both stimuli were presented, participants were cued to compare their spatial frequencies and respond. A typical trial is shown in Figure 1d.

#### Experiment 2: Spatial Incongruence on Visuotactile Integration

Experiment 2 aimed to investigate the effect of spatial proximity on visuotactile integration of spatial frequency. It followed a similar 2AFC design to the bimodal, congruent spatial frequency condition in Experiment 1. However, in the test trials, the visual stimulus was displaced horizontally from the tactile stimulus, and the 3D model of the device was covered by a large 10m-wide virtual tabletop, eliminating any visual reference for the spatial location of the visual stimuli (see Figure 1c.).

### Data analysis and MLE model prediction

#### Data Cleaning

In both experiments, trials with response time (RT) exceeding two standard deviations above the mean (Exp 1: RT > 2.91 s, Exp 2: RT > 2.24 s) were considered anomalies and removed from the analysis. This exclusion aimed to remove trials in which participants were disrupted by external events or experienced lapses. The exclusion method (z-score method) was based on the recommendations of Berger and Kiefer (2021), which should minimise the bias introduced by outlier trials.

#### Psychometric function fitting

The cumulative Gaussian psychometric function was fitted for every participant under each condition. The mean (*µ* or point of subjective equality, PSE) and standard deviation (*σ*, slope) of the cumulative Gaussian function can be used to quantify the sensitivity and bias in the participant’s spatial frequency perception respectively. With a higher *σ* representing worse sensitivity, and PSE representing the estimated testing stimulus frequency at which it is perceived the same as the reference stimulus(0.400 cycles/mm).

#### MLE prediction

The MLE integration model proposes that the bimodal estimation of spatial frequency is a weighted average of the unimodal inputs, with weights determined by the reliability of each modality. Specifically, the weights are inversely proportional to the variance of the unimodal sensory inputs (*σ*^2^), such that more reliable cues contribute more to the combined estimate:

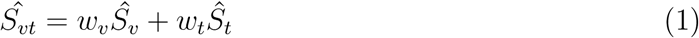

Where *Ŝ* represents the perceived stimulus and *w* represents the weights, which can be calculated as:

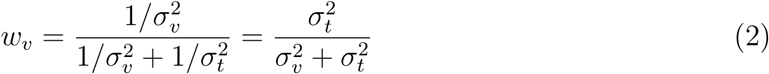

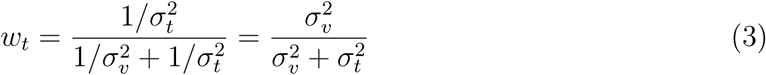

The MLE integration model also predicts that the variance for the bi-modal condition

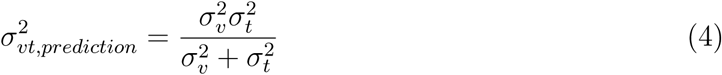

For the incongruent bi-modal condition, the PSE can also be predicted based on the MLE model. AS PSE denoted where the test and reference stimulus were perceived as equal (*Ŝ_vt,ref_* = *Ŝ_vt,test_*), hence we could get:

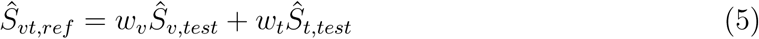

In the incongruent bi-modal condition, the spatial frequency of each test stimulus pair in the visual modality was always either lower or higher by Δ*f* in spatial frequency compared to touch, hence for the incongruent condition where vision had a lower frequency, we assume, for simplicity, that unimodal estimates are unbiased and that sensory integration follows linearity in the frequency space, we could then get

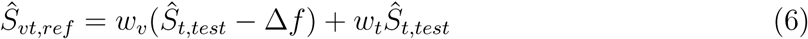

And:

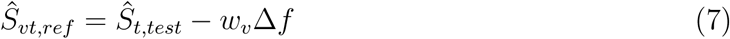

Hence, the PSE for incongruent, vision lower frequency condition, defined as the tactile test stimulus at which the perceived bimodal estimate is equal to the reference stimulus, can be calculated using the MLE model as follows:

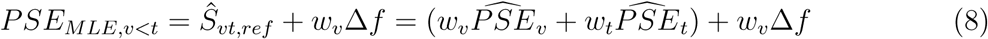

Similarly, for incongruent, vision higher frequency condition, we could get:

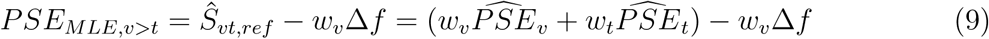

And for congruent bi-modal conditions:

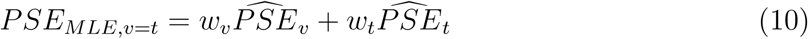

## Results

### Experiment 1: Visuotactile Integration of Surface Spatial Frequency

Experiment 1 aims to investigate whether the brain integrates spatial frequency information from vision and touch in a statistically optimal manner. Participants were tested under five different congruency conditions: visual-only, touch-only, congruent bi-modal, incongruent (visual lower frequency), and incongruent (visual higher frequency). Maximum likelihood estimation (MLE) model predictions were computed for each participant based on their uni-modal performance. These predictions included both spatial frequency precision (*σ*) for the biases (*PSE*) for bi-modal conditions (see the Methods section for further details).

### Effect of congruency condition on spatial frequency precision

A repeated-measures analysis of variance (RMANOVA) was conducted to test the main effect of congruency conditions on spatial frequency acuity (*σ*). Mauchly’s test indicated that the assumption of sphericity was violated (*χ*^2^(14) = 63.053, *p < .*001). Therefore, a Greenhouse-Geisser correction was applied. The main effect of congruency condition was significant after correction (*F* (2.182, 39.283) = 18.113, *p < .*001, 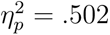).

*Post hoc* pairwise comparisons using the Holm-Bonferroni correction revealed that spatial frequency precision was significantly better for all three bi-modal conditions (congruent bi-modal, incongruent with visual lower frequency, and incongruent with visual higher frequency) as well as the MLE prediction, compared to either of the uni-modal conditions (vision-only, touch-only). No other significant pairwise comparisons were found (see Table 1 for detailed statistics). See Figure 3a and b for boxplot of empirical and MLE predicted *σ*.

**Figure 3.**
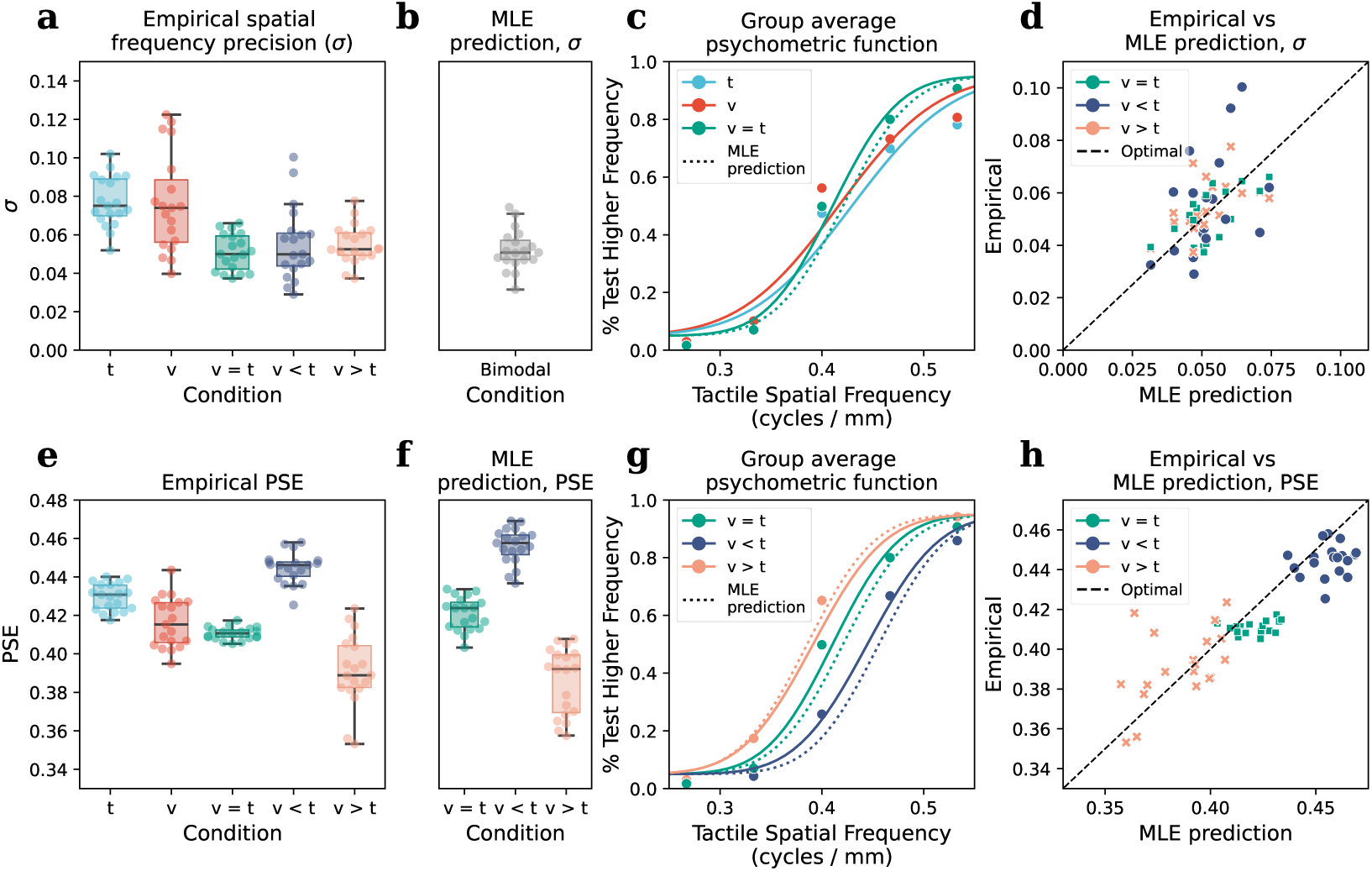
Experiment 1: Visuotactile integration of spatial frequency. *Note.* **a.** Boxplot of spatial frequency precision (*σ*) across all empirical condition in Experiment 1. *Post hoc* comparisons indicate that all three bimodal conditions exhibit better precision than either of the two unimodal conditions. **b.** MLE prediction of bimodal spatial frequency precision. No significant differences were found between the MLE predictions and the three bimodal conditions. **c.** Group-averaged psychometric functions for the two unimodal conditions and the congruent bimodal condition, along with the MLE-predicted psychometric function for the congruent bimodal condition (dotted line). The congruent bimodal condition and the MLE prediction exhibit greater precision than the unimodal conditions, as indicated by the steeper slopes. **d.** Empirical *σ* were plotted against MLE prediction, and a significant relationship was found between the two. No significant effects were found between different bi-modal conditions. **e.** Boxplot of PSE across all empirical conditions. **f.** MLE prediciton of PSE. **g.** Group-averaged psychometric functions for the three empirical bi-modal conditions and corresponding MLE prediction. **h.** Empirial PSE against MLE prediction. Both MLE prediction and congruency conditions were found to predict the empirical values significantly.

**Table 1.**
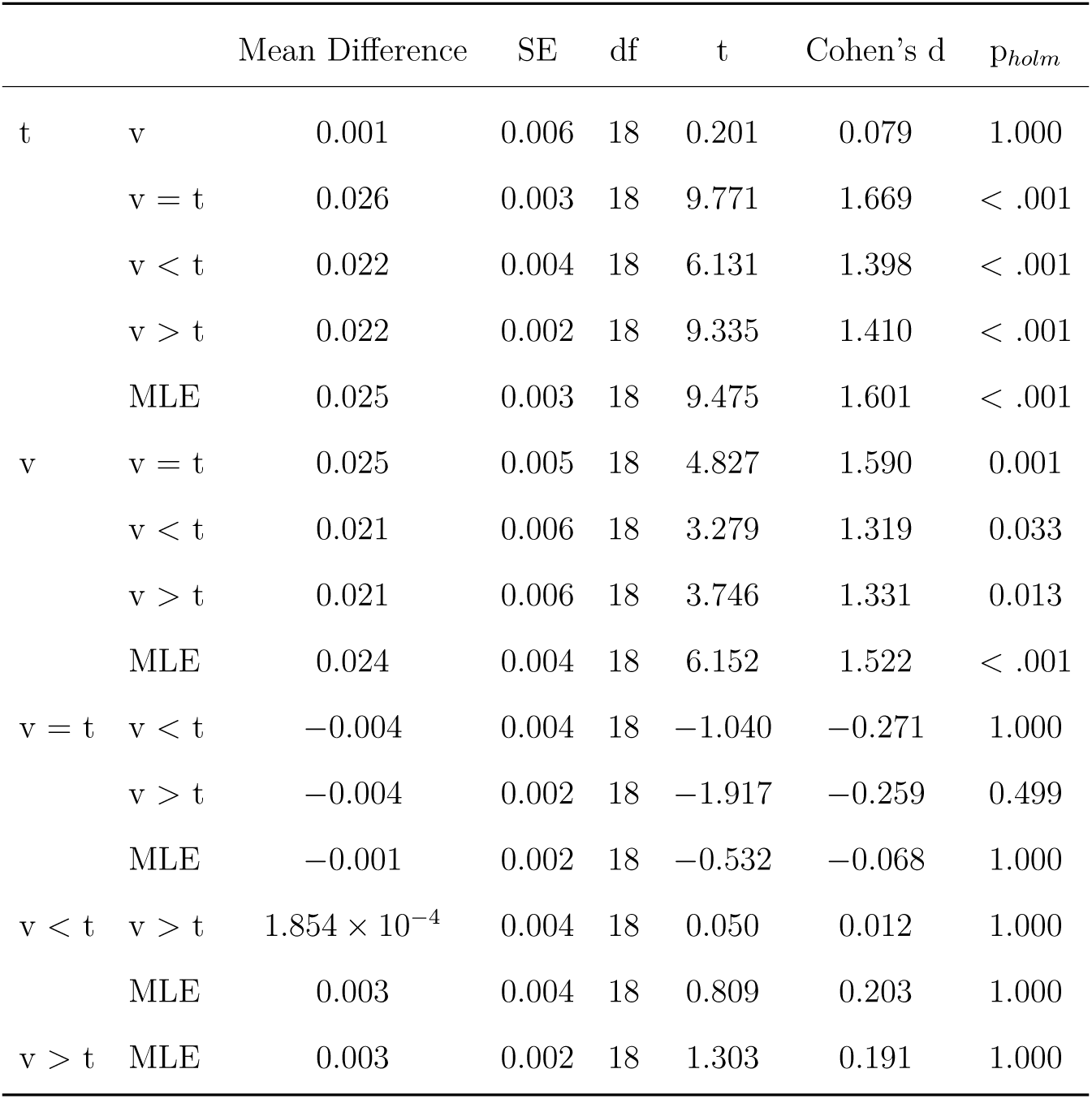
Experiment 1: Post Hoc Comparisons of Spatial Frequency Precision Across Frequency Congruency Conditions.

In addition to the RMANOVA, A multiple linear regression was conducted to examine whether the MLE prediction and bimodal congruency conditions significantly predict the empirical data (Figure 3d). For the regression analysis, bimodal congruency conditions (*v* = *t*, *v < t*, *v > t*) were coded as the difference between visual and tactile spatial frequencies (0, -0.067, 0.067 cycles/mm respectively) such that it is a continuous variable. The overall regression model was significant, *R*^2^ = 0.224, *F* (2, 54) = 7.798, *p* = 0.001, indicating that MLE prediction and congruency conditions explain a significant portion of the variance in the empirical data. The regression coefficients revealed that MLE prediction was a significant predictor of empirical *σ* (*b* = 0.624, *β* = 0.473, *SE* = 0.157, *t*(54) = 3.949, *p < .*001). However, the congruency condition was not a significant predictor of empirical spatial frequency precision (*σ*) (*b* = *−*0.001, *β* = *−*0.006, *SE* = 0.030, *t*(54) = *−*0.047, *p* = .963).

These findings strongly support the hypothesis of multisensory integration of spatial frequency information between vision and touch. All bimodal conditions exhibited significantly better precision than either unimodal condition, highlighting the facilitative effect of combining multisensory cues. Furthermore, the lack of significant differences between the MLE predictions and the three bimodal conditions suggests that the experimental results align well with the MLE model. Additionally, regression analysis indicated that congruency had no effect on spatial frequency precision, further reinforcing the MLE integration hypothesis.

### Effect of congruency condition on biases in spatial frequency perception

To further test whether the incongruence of tactile and visual stimulus would lead to biases in spatial frequency perception, and whether the biases (PSE) can be predicted with the MLE model, another RMANOVA was conducted to test the main effect of congruency conditions on PSE. As the predicted PSEs were MLE predictions based on linear combinations of empirical PSEs, Mauchly’s test cannot be conducted due to collinearity between conditions. Instead, John, Nagao and Sugiura (JNS) test of sphericity (Abdi, 2010) was conducted and indicated that the assumption of sphericity was violated 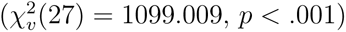. And a Greenhouse-Geisser correction was applied. The main effect of congruency condition was significant after correction (*F* (2.506, 45.108) = 102.853, *p < .*001, 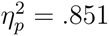).

Figure 3e and f show the boxplots for PSE across all conditions, and the statistics for the *post hoc* comparisons are detailed in Table 2. Several key comparisons are as follows: The incongruent, vision-lower-frequency condition was found to have a PSE larger than all other empirical conditions, while the incongruent, vision-higher-frequency condition had a PSE lower than all other conditions. The PSE for the empirical congruent bimodal condition was significantly higher than the MLE prediction. No significant difference was found between the incongruent, vision-lower-frequency condition and its MLE prediction, which confirms the MLE prediction. Additionally, the empirical incongruent, vision-higher-frequency condition had a PSE significantly higher than its MLE prediction.

**Table 2.** Experiment 1: Post Hoc Comparisons of PSE Across Spatial Frequency Congruency Conditions.

| | | Mean Difference | SE | df | t | Cohen's d | $p_{holm}$ |
| --- | --- | --- | --- | --- | --- | --- | --- |
| t | v | 0.013 | 0.003 | 18 | 3.857 | 1.189 | 0.005 |
|  | v = t | 0.019 | 0.002 | 18 | 11.889 | 1.716 | < .001 |
|  | v < t | -0.015 | 0.002 | 18 | -7.426 | -1.375 | < .001 |
|  | v > t | 0.038 | 0.004 | 18 | 8.497 | 3.453 | < .001 |
|  | v = t, MLE | 0.009 | 0.002 | 18 | 4.293 | 0.787 | 0.003 |
|  | v < t, MLE | -0.016 | 0.002 | 18 | -7.022 | -1.475 | < .001 |
|  | v > t, MLE | 0.054 | 0.003 | 18 | 15.533 | 4.907 | < .001 |
| v | v = t | 0.006 | 0.003 | 18 | 1.895 | 0.527 | 0.149 |
|  | v < t | -0.028 | 0.004 | 18 | -8.005 | -2.565 | < .001 |
|  | v > t | 0.025 | 0.004 | 18 | 6.358 | 2.263 | < .001 |
|  | v = t, MLE | -0.004 | 0.002 | 18 | -2.666 | -0.402 | 0.047 |
|  | v < t, MLE | -0.029 | 0.005 | 18 | -5.818 | -2.665 | < .001 |
|  | v > t, MLE | 0.041 | 0.002 | 18 | 17.144 | 3.718 | < .001 |
| v = t | v < t | -0.034 | 0.002 | 18 | -17.662 | -3.092 | < .001 |
|  | v > t | 0.019 | 0.004 | 18 | 4.416 | 1.737 | 0.002 |
|  | v = t, MLE | -0.010 | 0.002 | 18 | -5.274 | -0.929 | < .001 |
|  | v < t, MLE | -0.035 | 0.002 | 18 | -14.113 | -3.191 | < .001 |
|  | v > t, MLE | 0.035 | 0.002 | 18 | 14.113 | 3.191 | < .001 |
| v < t | v > t | 0.053 | 0.004 | 18 | 12.189 | 4.828 | < .001 |
|  | v = t, MLE | 0.024 | 0.003 | 18 | 9.325 | 2.163 | < .001 |
|  | v < t, MLE | -0.001 | 0.003 | 18 | -0.362 | -0.100 | 0.722 |
|  | v > t, MLE | 0.069 | 0.003 | 18 | 21.222 | 6.283 | < .001 |
| v > t | v = t, MLE | -0.029 | 0.004 | 18 | -7.439 | -2.666 | < .001 |
|  | v < t, MLE | -0.054 | 0.006 | 18 | -9.136 | -4.928 | < .001 |
|  | v > t, MLE | 0.016 | 0.004 | 18 | 4.186 | 1.454 | 0.003 |
| v = t, MLE | v < t, MLE | -0.025 | 0.004 | 18 | -6.372 | -2.263 | < .001 |
|  | v > t, MLE | 0.045 | 0.002 | 18 | 21.081 | 4.120 | < .001 |
| v < t, MLE | v > t, MLE | 0.070 | 0.005 | 18 | 14.113 | 6.382 | < .001 |

The analysis of biases (PSEs) in spatial frequency perception also supports the hypothesis of MLE integration of spatial frequency cues between touch and vision, as the PSEs for incongruent conditions were found to shift according to the incongruent visual stimulus. However, for the incongruent, vision-higher-frequency and congruent bimodal conditions, the MLE model appears to overestimate the biases. The implication of this particular finding will be discussed in detail in the Discussion section.

A multiple linear regression analysis was also conducted for PSE, and the overall regression model was significant (*R*^2^ = 0.802, *F* (2, 54) = 109.323, *p <* 0.001). The regression coefficients revealed that both MLE prediction (*b* = 0.393, *β* = 0.124, *SE* = 0.124, *t*(54) = 3.158, *p* = .003) and congruency condition (*b* = *−*0.191, *β* = *−*0.420, *SE* = 0.071, *t*(54) = *−*2.693, *p* = .009) significantly predicted the empirical PSE.

The results of the regression analysis on PSE also support the MLE integration model, as the MLE prediction significantly predicted the empirical PSEs in all bimodal conditions. Additionally, the empirical PSE was significantly shifted by the incongruent visual frequency, complementing the findings from the previous RMANOVA on PSE.

### Effect of congruency condition on response time

To better understand the difference in sensory processing between each congruency condition, an RMANOVA was performed to test the effect of congruency condition on response time. Mauchly’s test indicated that the assumption of sphericity was violated (*χ*^2^(9) = 28.519, *p < .*001), and a Greenhouse-Geisser correction was applied. The main effect of congruency condition was significant after correction (*F* (2.470, 44.465) = 35.081, *p < .*001, 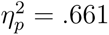). indicating that response times varied across conditions.

*Post hoc* comparisons using the Holm-Bonferroni correction showed that response times in the touch-only condition were significantly slower than all other conditions, and the vision-only conditions were significantly faster than all other conditions. No significant difference was found in response time between the three bimodal conditions. See Table 3 for detailed statistics.

**Table 3.** Experiment 1: Post Hoc Comparisons of Response Time Across Frequency Congruency Conditions.

| | | Mean Difference | SE | df | t | Cohen's d | $p_{holm}$ |
| --- | --- | --- | --- | --- | --- | --- | --- |
| t | v | 0.308 | 0.040 | 18 | 7.627 | 1.205 | < .001 |
|  | v = t | 0.113 | 0.025 | 18 | 4.447 | 0.442 | 0.002 |
|  | v < t | 0.083 | 0.028 | 18 | 2.929 | 0.326 | 0.038 |
|  | v > t | 0.079 | 0.026 | 18 | 3.001 | 0.309 | 0.038 |
| v | v = t | -0.195 | 0.025 | 18 | -7.868 | -0.762 | < .001 |
|  | v < t | -0.224 | 0.033 | 18 | -6.861 | -0.878 | < .001 |
|  | v > t | -0.229 | 0.031 | 18 | -7.414 | -0.896 | < .001 |
| v = t | v < t | -0.030 | 0.024 | 18 | -1.230 | -0.116 | 0.469 |
|  | v > t | -0.034 | 0.015 | 18 | -2.250 | -0.133 | 0.112 |
| v < t | v > t | -0.004 | 0.017 | 18 | -0.263 | -0.017 | 0.795 |

### Experiment 2: Effect of spatial proximity on visuotactile integration of spatial frequency cue

Experiment 2 aims to investigate whether spatial proximity affects the integration of spatial frequency cues between vision and touch. This experiment consists only of bimodal trials with congruent visual and tactile spatial frequencies. However, the visual stimulus was horizontally displaced from the tactile stimulus at five different distances to examine whether spatial proximity influences the integration of spatial frequency cues from the two modalities.

### Effect of spatial proximity on spatial frequency precision

A repeated-measures ANOVA was conducted to test the main effect of spatial proximity on spatial frequency precision (*σ*). Mauchly’s test indicated that the assumption of sphericity was violated (*χ*^2^(9) = 38.096, *p < .*001). Therefore, a Greenhouse-Geisser correction was applied. The main effect of spatial proximity was not significant after correction (*F* (1.810, 32.578) = 1.973, *p* = .159, 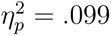). To further understand whether the spatial frequency precision of the bimodal perception was affected by spatial proximity, a Bayesian RMANOVA was performed, and the Bayes factor indicated anecdotal evidence in favour of the null hypothesis, *BF*_10_ = 0.543. Figure 5a shows the boxplot of spatial frequency precision (*σ*) at different horizontal displacement levels.

### Effect of spatial proximity on biases in spatial frequency perception

Another repeated-measures ANOVA was conducted to test the effect of spatial proximity on biases (PSE) in spatial frequency perception. Mauchly’s test indicated that the assumption of sphericity was violated (*χ*^2^(9) = 27.261, *p* = .001). The main effect of spatial proximity on PSE after Greenhouse-Geisser correction is significant (*F* (2.693, 48.477) = 3.196, *p* = .036, 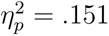). However, *post hoc* tests with Holm-Bonferroni correction find no significant difference between any pair of displacement conditions (see Table 4). The Bayes factor indicated anecdotal evidence in favour of the alternative hypothesis, *BF*_10_ = 0.543.

**Table 4.** Experiment 2: Post hoc comparisons of PSE across horizontal displacement conditions.

| | | Mean Difference | SE | df | t | $p_{holm}$ |
| --- | --- | --- | --- | --- | --- | --- |
| 0 | 3 cm | 0.006 | 0.003 | 18 | 2.472 | 0.189 |
|  | 6 cm | 0.018 | 0.007 | 18 | 2.725 | 0.125 |
|  | 9 cm | 0.013 | 0.007 | 18 | 1.879 | 0.536 |
|  | 12 cm | 0.014 | 0.005 | 18 | 2.987 | 0.079 |
| 3 cm | 6 cm | 0.011 | 0.006 | 18 | 1.845 | 0.536 |
|  | 9 cm | 0.006 | 0.005 | 18 | 1.180 | 1.000 |
|  | 12 cm | 0.008 | 0.004 | 18 | 1.801 | 0.536 |
| 6 cm | 9 cm | -0.005 | 0.007 | 18 | -0.697 | 1.000 |
|  | 12 cm | -0.004 | 0.005 | 18 | -0.697 | 1.000 |
| 9 cm | 12 cm | 0.001 | 0.005 | 18 | 0.250 | 1.000 |

### Effect of spatial proximity on response time

Similar to Experiment 1, an RMANOVA was conducted to examine the effect of spatial proximity on response time. Mauchly’s test indicated that the assumption of sphericity was not violated (*χ*^2^(9) = 5.213*, p* = .816). No significant main effect of displacement on response time was found (*F* (4, 72) = 0.133, *p* = .970, 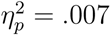).

Overall, despite the inconclusive results in PSE analysis, the analysis of spatial frequency precision and biases found no clear evidence that spatial proximity affects the degree of visuotactile integration of spatial frequency cues. Furthermore, the absence of differences in response time across displacement conditions provides additional support for the notion that spatial proximity does not influence the sensory processing of bimodal information.

## Discussion

### Visuo-tactile integration of spatial frequency cue follows optimal integration

In Experiment 1, we demonstrated that spatial integration between vision and touch follows the MLE rule, with bimodal conditions showing better precision than unimodal conditions. The MLE model successfully predicted the improvement in spatial frequency precision under the bimodal condition. Additionally, the PSE analysis revealed that when incongruent visual and tactile frequencies were presented, the bimodal perception fell between the individual unimodal perceptions. Regression analysis on spatial frequency precision and bias also finds that MLE prediction significantly accounts for the empirical data, which also supports the model.

One interesting finding that seems to challenge the MLE model is the PSE prediction for incongruent, vision lower frequency condition and congruent bi-modal condition, the RMANOVA showed that the MLE prediction in shift is more extreme than the actual PSE shift, which suggests that the weight of visual modality is lower than the MLE prediction in the lower frequency range. One possible explanation for these findings is the nonlinearity of spatial frequency space.

Nonlinear behaviour in contrast sensitivity is well-documented in the literature, with human contrast sensitivity varying nonlinearly across different spatial frequency ranges (Campbell & Robson, 1968; Hall & Hall, 1977; Pointer & Hess, 1989). Similarly, in somatosensory perception, substantial evidence suggests that tactile spatial sensitivity varies nonlinearly across different frequency ranges, with an optimal range of spatial frequencies at which sensitivity is maximised (Hsiao et al., 1993). Besides, there were also various studies that suggest temporal cues can influence tactile spatial acuity (Cascio & Sathian, 2001; Gamzu & Ahissar, 2001). In an active exploration setting like the current experiment, changes in the spatial frequency of surface texture are likely to alter the rate of temporal cues received by mechanoreceptive afferents in the glabrous skin of the fingertip as the finger moves across the tactile stimulus. Previous studies have demonstrated nonlinear behaviour in tactile spatial acuity across different vibratory frequencies (Bensmaïa et al., 2006) and tactile spatial perception, such as orientation acuity, could be influenced by the introduction of such temporal cue during active exploration (Wang & Alais, 2025). Therefore, variability in temporal information may also contribute to the overall nonlinearity observed across different frequency ranges.

In Experiment 1, we also found evidence of nonlinearity in tactile and visual frequency perception, with significant differences in PSE between vision-only, tactile-only, and congruent bimodal conditions (see Figure 3e and Table 4). The difference in PSE between the two unimodal conditions and the congruent bimodal conditions suggests that, firstly, there might be nonlinearity in sensitivity at different frequencies, and secondly, the sensitivity profile might also differ between the two modalities. The nonlinear sensitivity across the spatial frequency space implies that each modality’s weight may change across frequency ranges, depending on its reliability at a given spatial frequency, rather than remaining constant. This nonlinearity could explain the overestimation observed in the MLE model for congruent bimodal and incongruent higher frequency visual conditions.

Another possibility for the observed PSE being less affected by vision at vision lower frequency condition compared to the model prediction is the influence of the central tendency effect, as suggested by Aston et al. (2021). In the experimental design, incongruent trials were interleaved with the other three conditions (vision-only, touch-only, and congruent bimodal). The trial sequence was pseudorandomised to ensure an equal number of trials for each condition within each experimental block. Additionally, the average of all trials in a block was centred around the reference spatial frequency (0.4 cycles/mm), with more than half of the visual and tactile stimuli (including all reference stimuli and part of the testing stimuli) also set at 0.4 cycles/mm. As a result, participants’ perceptions and decision-making may have been biased toward the average stimulus. This could explain the observed difference between the MLE prediction and the empirical PSE for the incongruent conditions. Even though the integration of incongruent vision and tactile cues drives bimodal perception away from the tactile stimulus and towards the visual stimulus (which has a higher or lower spatial frequency), the central tendency effect would also pull the bimodal perception and decision towards the average stimulus (which equals the reference stimulus of 0.4 cycles/mm). This, in turn, could reduce the shift in the PSE. In the Bayesian framework, this central tendency could also be viewed as a prior distribution with its mean at the reference stimulus, thereby influencing the posterior distribution of the bimodal perception accordingly.

### Optimal integration of spatial frequency cue is not affected by spatial proximity

Unlike the findings of Gepshtein et al. (2005), our results from Experiment 2 found no evidence that supports the hypothesis that spatial proximity affects integration. This discrepancy may arise from several factors, offering a valuable opportunity to explore the underlying mechanisms of multisensory cue integration.

One key factor that might lead to the difference between the two studies is the tasks and features tested. As Spence (2013) suggested, spatial proximity effects are less likely when tasks do not require spatial location information. In Gepshtein et al. (2005)’s study, the integrated feature was inter-surface distance, which required participants to focus on the location of surfaces, and engaging spatial attention. In contrast, our study examined spatial frequency perception, which is a localised feature that does not inherently depend on spatial location information, this could explain the absence of a spatial proximity effect.

Besides, Takahashi et al. (2009) also found that the brain integrates visual and tactile signals in an optimal fashion during tool use, even when there is a large spatial offset between the visual signal of the object at the tool tip and the tactile signal from the hand. However, this integration was reduced when the visual signal was moved away from the tool tip. This finding provides strong evidence suggesting that spatial proximity itself might not be the determining factor in whether information from different modalities is integrated. Instead, spatial proximity appears to provide information that helps the brain determine whether cues from different modalities originate from the same source.

Complementing this hypothesis, this benefit of common source in multisensory integration between vision and touch has been demonstrated by Helbig and Ernst (2007a).

Another consideration that further supports this hypothesis is that different integration mechanisms may be recruited depending on the nature of the task. Sathian and Zangaladze (2002) demonstrated that distinct patterns of brain activation occur when participants attend to different features within the same tactile stimulus. Specifically, orientation processing engaged the visual cortex, whereas spatial frequency perception did not. Similarly, orientation also requires awareness and reference to the location of the stimulus, where spatial frequency perception does not. Complementing this, Stoesz et al. (2003) also find macrospatial features (large scale features like shape, size and orientation) elicit different brain activity when compared to microspatial features (smaller, localised features like texture). Together, these findings suggest that there might be differential deployment of the visual system in tactile tasks.

Another interesting finding that seems to support the hypothesis that the integration of visual and tactile spatial frequency occurs at later stages comes from the discrepancy in response times across different congruency conditions in Experiment 1. Previous research has shown that human observers can detect combinations of multisensory signals more quickly than when each signal is presented individually (Hecht et al., 2008).

However, in Experiment 1, we observed a different pattern: the vision-only condition was consistently faster in response time compared to the bimodal conditions, while the tactile-only condition was consistently slower (see Figure 4a). This finding seems to suggest that for spatial frequency information, unimodal information may first be processed independently and only combined at higher cognitive hierarchical levels. This could explain the slower yet more precise bimodal perception compared to the vision-only condition.

**Figure 4.**
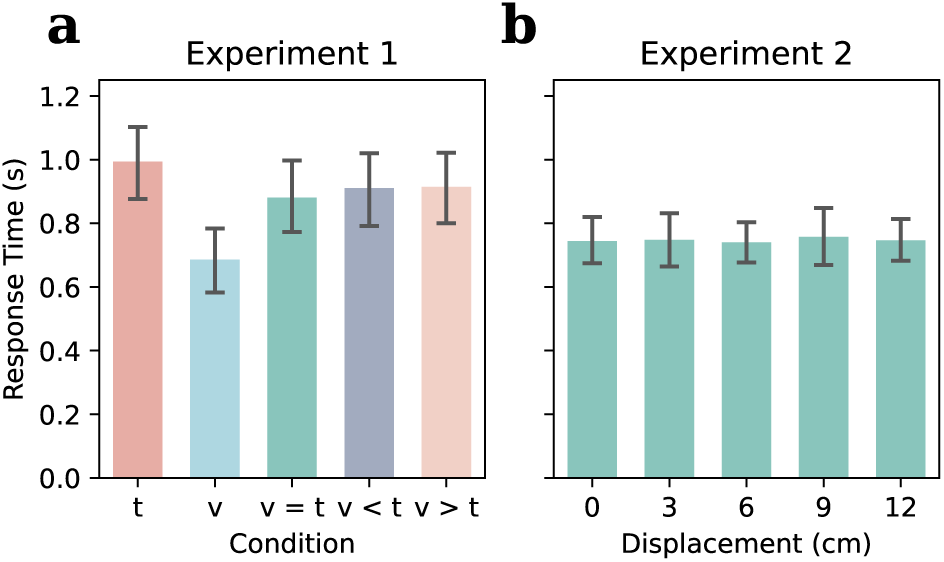
Experiment 1: Effect of congruency condition on response time. *Note.* **a.** Bar plot of mean response time (RT) across congruency conditions in Experiment 1. The tactile-only condition had significantly longer RTs compared to all other conditions, while the vision-only condition had the shortest RTs. **b.** Bar plot of mean RT across congruency conditions in Experiment 2. No significant main effect of spatial proximity on RT was found.

**Figure 5.**
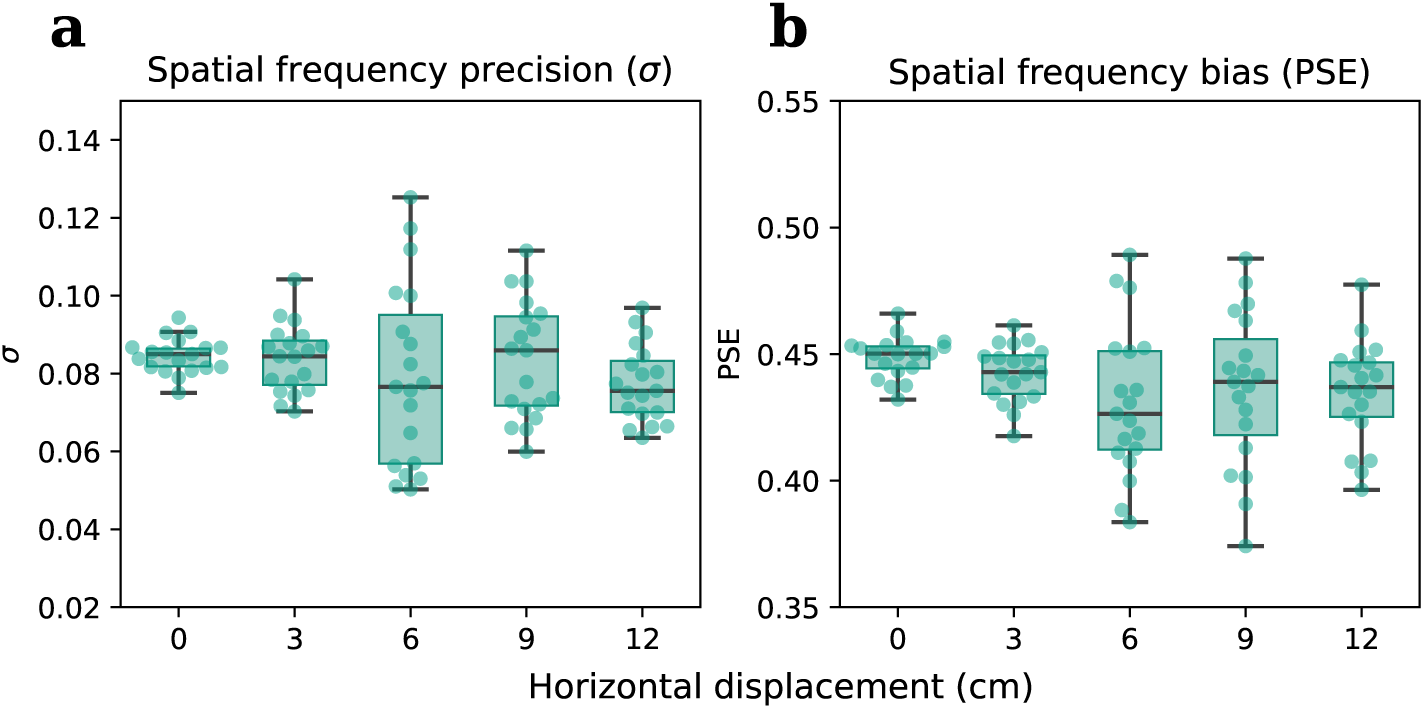
Experiment 2: Effect of spatial proximity on visuotactile integration of spatial frequency. *Note.* **a.** Box plot of spatial frequency precision (*σ*) across five different horizontal displacements. No significant effect of spatial proximity on spatial frequency precision was found. **b.** Box plot of biases in spatial frequency perception (PSE) across five different horizontal displacements. A significant main effect was found, but no pairwise comparisons were significant.

Furthermore, the independent processing of unimodal information before integration may also account for the lack of effect of spatial proximity on integration. The facilitative effect of multisensory integration might arise from higher-level cognitive processes, where the information is more abstract and not tied to a specific receptive field location. The response time results in Experiment 2 also support this, as no significant difference was found in response times across different displacements (Figure 4b). This also seems to suggest that similar integration processes occur despite the discrepancy in spatial locations, with no evidence of an additional facilitative effect when the stimuli were colocalised.

Appendix

## Notes

### Competing Interest Statement

The authors have declared no competing interest.

### Funding

- Australian Research Council: DP210101691

## References

Abdi, H. (2010). The greenhouse-geisser correction. In The greenhouse.

Alais, D., & Burr, D. (2004). The ventriloquist effect results from near-optimal bimodal integration [Publisher: Elsevier]. Current Biology, 14 (3), 257–262. 10.1016/j.cub.2004.01.029

Alais, D., & Burr, D. (2019). Cue combination within a bayesian framework. In A. K. C. Lee, M. T. Wallace, A. B. Coffin, A. N. Popper, & R. R. Fay (Eds.), Multisensory processes: The auditory perspective (pp. 9–31). Springer International Publishing. 10.1007/978-3-030-10461-0_2

Arnold, D. H., Petrie, K., Murray, C., & Johnston, A. (2019). Suboptimal human multisensory cue combination [Publisher: Nature Publishing Group]. Scientific Reports, 9 (1), 5155. 10.1038/s41598-018-37888-7

Aston, S., Negen, J., Nardini, M., & Beierholm, U. (2021). Central tendency biases must be accounted for to consistently capture bayesian cue combination in continuous response data. Behavior Research Methods, 54 (1), 508–521. 10.3758/s13428-021-01633-2

Bensmaïa, S. J., Craig, J. C., & Johnson, K. O. (2006). Temporal factors in tactile spatial acuity: Evidence for RA interference in fine spatial processing [Publisher: American Physiological Society]. Journal of Neurophysiology, 95 (3), 1783–1791. 10.1152/jn.00878.2005

Berger, A., & Kiefer, M. (2021). Comparison of different response time outlier exclusion methods: A simulation study [Publisher: Frontiers]. Frontiers in Psychology, 12. 10.3389/fpsyg.2021.675558

Bourgeon, S., Dépeault, A., Meftah, E.-M., & Chapman, C. E. (2016). Tactile texture signals in primate primary somatosensory cortex and their relation to subjective roughness intensity [Publisher: American Physiological Society]. Journal of Neurophysiology, 115 (4), 1767–1785. 10.1152/jn.00303.2015

Campbell, F. W., & Robson, J. G. (1968). Application of fourier analysis to the visibility of gratings. The Journal of Physiology, 197 (3), 551–566. Retrieved March 25, 2025, from https://www.ncbi.nlm.nih.gov/pmc/articles/PMC1351748/

Cascio, C. J., & Sathian, K. (2001). Temporal cues contribute to tactile perception of roughness [Publisher: Society for Neuroscience Section: ARTICLE]. The Journal of Neuroscience, 21 (14), 5289–5296. 10.1523/JNEUROSCI.21-14-05289.2001

De Valois, R. L., Albrecht, D. G., & Thorell, L. G. (1982). Spatial frequency selectivity of cells in macaque visual cortex. Vision Research, 22 (5), 545–559. 10.1016/0042-6989(82)90113-4

DeValois, R. L., & DeValois, K. K. (1990). Spatial vision. Oxford University Press, Incorporated. https://books.google.com.au/books?id=%5C_fO6wzoVK2wC

Ernst, M. O., & Banks, M. S. (2002). Humans integrate visual and haptic information in a statistically optimal fashion [Publisher: Nature Publishing Group UK London]. Nature, 415 (6870), 429–433. 10.1038/415429a

Fetsch, C. R., DeAngelis, G. C., & Angelaki, D. E. (2010). Visual–vestibular cue integration for heading perception: Applications of optimal cue integration theory [_eprint: https://onlinelibrary.wiley.com/doi/pdf/10.1111/j.1460-9568.2010.07207.x]. *European Journal of Neuroscience*, *31* (10), 1721–1729. 10.1111/j.1460-9568.2010.07207.x

Foster, K. H., Gaska, J. P., Nagler, M., & Pollen, D. A. (1985). Spatial and temporal frequency selectivity of neurones in visual cortical areas v1 and v2 of the macaque monkey. [_eprint: https://onlinelibrary.wiley.com/doi/pdf/10.1113/jphysiol.1985.sp015776]. *The Journal of Physiology*, *365* (1), 331–363. 10.1113/jphysiol.1985.sp015776

Gamzu, E., & Ahissar, E. (2001). Importance of temporal cues for tactile spatial- frequency discrimination [Publisher: Society for Neuroscience Section: ARTICLE]. The Journal of Neuroscience, 21 (18), 7416–7427. 10.1523/JNEUROSCI.21-18-07416.2001

Gepshtein, S., Burge, J., Ernst, M. O., & Banks, M. S. (2005). The combination of vision and touch depends on spatial proximity [Publisher: Association for Research in Vision and Ophthalmology (ARVO)]. Journal of Vision, 5 (11), 7. 10.1167/5.11.7

Guest, S., & Spence, C. (2003). What role does multisensory integration play in the visuotactile perception of texture? International Journal of Psychophysiology, 50 (1), 63–80. 10.1016/S0167-8760(03)00125-9

Hall, C. F., & Hall, E. L. (1977). A nonlinear model for the spatial characteristics of the human visual system [Conference Name: IEEE Transactions on Systems, Man, and Cybernetics]. *IEEE Transactions on Systems*, Man, and Cybernetics, 7 (3), 161–170. 10.1109/TSMC.1977.4309680

Hecht, D., Reiner, M., & Karni, A. (2008). Multisensory enhancement: Gains in choice and in simple response times. Experimental Brain Research, 189 (2), 133–143. 10.1007/s00221-008-1410-0

Helbig, H. B., & Ernst, M. O. (2007a). Knowledge about a common source can promote visual — haptic integration [Publisher: SAGE Publications Ltd STM]. Perception, 36 (10), 1523–1533. 10.1068/p5851

Helbig, H. B., & Ernst, M. O. (2007b). Optimal integration of shape information from vision and touch. Experimental Brain Research, 179 (4), 595–606. 10.1007/s00221-006-0814-y

Helbig, H. B., Ernst, M. O., Ricciardi, E., Pietrini, P., Thielscher, A., Mayer, K. M., Schultz, J., & Noppeney, U. (2012). The neural mechanisms of reliability weighted integration of shape information from vision and touch [Publisher: Elsevier BV]. NeuroImage, *60* (2), 1063–1072. 10.1016/j.neuroimage.2011.09.072

Hsiao, S. S., Johnson, K. O., & Twombly, I. (1993). Roughness coding in the somatosensory system. Acta Psychologica, 84 (1), 53–67. 10.1016/0001-6918(93)90072-Y

Jones, B., & O’Neil, S. (1985). Combining vision and touch in texture perception. Perception & Psychophysics, 37 (1), 66–72. 10.3758/BF03207140

Kadunce, D. C., Vaughan, W. J., Wallace, M. T., & Stein, B. E. (2001). The influence of visual and auditory receptive field organization on multisensory integration in the superior colliculus. Experimental Brain Research, 139 (3), 303–310. 10.1007/s002210100772

Kauffmann, L., Ramanoël, S., & Peyrin, C. (2014). The neural bases of spatial frequency processing during scene perception [Publisher: Frontiers]. Frontiers in Integrative Neuroscience, 8. 10.3389/fnint.2014.00037

Kelly, D. (1975). Spatial frequency selectivity in the retina. Vision Research, 15 (6), 665–672. 10.1016/0042-6989(75)90282-5

Lamb, M. R., & Yund, E. W. (1996). Spatial frequency and attention: Effects of level-, target-, and location-repetition on the processing of global and local forms. Perception & Psychophysics, 58 (3), 363–373. 10.3758/BF03206812

Lederman, S. J., Thorne, G., & Jones, B. (1986). Perception of texture by vision and touch: Multidimensionality and intersensory integration. [Place: US Publisher: American Psychological Association]. Journal of Experimental Psychology: Human Perception and Performance, 12 (2), 169–180. 10.1037/0096-1523.12.2.169

Lunghi, C., & Alais, D. (2013). Touch interacts with vision during binocular rivalry with a tight orientation tuning [Publisher: Public Library of Science San Francisco, USA]. PLoS ONE, 8 (3), e58754. 10.1371/journal.pone.0058754

Meijer, D., Veselič, S., Calafiore, C., & Noppeney, U. (2019). Integration of audiovisual spatial signals is not consistent with maximum likelihood estimation. Cortex, 119, 74–88. 10.1016/j.cortex.2019.03.026

Nikbakht, N., Tafreshiha, A., Zoccolan, D., & Diamond, M. E. (2018). Supralinear and supramodal integration of visual and tactile signals in rats: Psychophysics and neuronal mechanisms. Neuron, 97 (3), 626–639.e8. 10.1016/j.neuron.2018.01.003

Pointer, J., & Hess, R. (1989). The contrast sensitivity gradient across the human visual field: With emphasis on the low spatial frequency range. Vision Research, 29 (9), 1133–1151. 10.1016/0042-6989(89)90061-8

Pollen, D. A., & Ronner, S. F. (1983). Visual cortical neurons as localized spatial frequency filters [Conference Name: IEEE Transactions on Systems, Man, and Cybernetics]. *IEEE Transactions on Systems, Man, and Cybernetics*, SMC*-*13 (5), 907–916. 10.1109/TSMC.1983.6313086

Roberts, R. D., Li, M., & Allen, H. A. (2024). Visual effects on tactile texture perception [Publisher: Nature Publishing Group]. Scientific Reports, 14 (1), 632. 10.1038/s41598-023-50596-1

Sachs, M. B., Nachmias, J., & Robson, J. G. (1971). Spatial-frequency channels in human vision* [Publisher: Optica Publishing Group]. JOSA, 61 (9), 1176–1186. 10.1364/JOSA.61.001176

Sathian, K., & Burton, H. (1991). The role of spatially selective attention in the tactile perception of texture. Perception & Psychophysics, 50 (3), 237–248. 10.3758/BF03206747

Sathian, K., & Zangaladze, A. (2002). Feeling with the mind’s eye: Contribution of visual cortex to tactile perception [Publisher: Elsevier]. Behavioural Brain Research, 135 (1), 127–132. 10.1016/S0166-4328(02)00141-9

Sathian, K., Zangaladze, A., Hoffman, J. M., & Grafton, S. T. (1997). Feeling with the mind’s eye [Publisher: LWW]. NeuroReport, 8 (18), 3877–3881. 10.1097/00001756-199712220-00008

Spence, C. (2013). Just how important is spatial coincidence to multisensory integration? evaluating the spatial rule. Annals of the New York Academy of Sciences, 1296 (1), 31–49. 10.1111/nyas.12121

Stein, B. E., Meredith, M. A., Huneycutt, W. S., & McDade, L. (1989). Behavioral indices of multisensory integration: Orientation to visual cues is affected by auditory stimuli. Journal of Cognitive Neuroscience, 1 (1), 12–24. 10.1162/jocn.1989.1.1.12

Stein, B. E., & Stanford, T. R. (2008). Multisensory integration: Current issues from the perspective of the single neuron [Publisher: Nature Publishing Group]. Nature Reviews Neuroscience, 9 (4), 255–266. 10.1038/nrn2331

Stoesz, M. R., Zhang, M., Weisser, V. D., Prather, S., Mao, H., & Sathian, K. (2003). Neural networks active during tactile form perception: Common and differential activity during macrospatial and microspatial tasks [Publisher: Elsevier]. International Journal of Psychophysiology, 50 (1), 41–49. 10.1016/S0167-8760(03)00123-5

Takahashi, C., Diedrichsen, J., & Watt, S. J. (2009). Integration of vision and haptics during tool use. Journal of Vision, 9 (6), 3. 10.1167/9.6.3

van der Groen, O., van der Burg, E., Lunghi, C., & Alais, D. (2013). Touch influences visual perception with a tight orientation-tuning [Publisher: Public Library of Science San Francisco, USA]. PLoS ONE, 8 (11), e79558. 10.1371/journal.pone.0079558

Wang, G., & Alais, D. (2024). Tactile adaptation to orientation produces a robust tilt aftereffect and exhibits crossmodal transfer when tested in vision [Publisher: Nature Publishing Group]. Scientific Reports, 14 (1), 10164. 10.1038/s41598-024-60343-9

Wang, G., & Alais, D. (2025, March 19). Characteristics of tactile orientation perception: Oblique effect, active vs passive exploration, and serial dependence [Pages: 2025.03.19.644099 Section: New Results]. 10.1101/2025.03.19.644099

Wang, G., Alais, D., Blake, R., & Han, S. (2022). CFS-crafter: An open-source tool for creating and analyzing images for continuous flash suppression experiments [Publisher: Springer]. Behavior Research Methods, 55 (4), 2004–2020. 10.3758/s13428-022-01903-7

Whitaker, T. A., Simões-Franklin, C., & Newell, F. N. (2008). Vision and touch: Independent or integrated systems for the perception of texture? Brain Research, 1242, 59–72. 10.1016/j.brainres.2008.05.037

Williams, D. W., Wilson, H. R., & Cowan, J. D. (1982). Localized effects of spatial-frequency adaptation [Publisher: Optica Publishing Group]. JOSA, 72 (7), 878–887. 10.1364/JOSA.72.000878

